# A functional perspective on the conditional covariance comparison problem in dementia analysis

**DOI:** 10.1101/2023.12.19.572366

**Authors:** Calvin Guan, Ashis Gangopadhyay, The Alzheimer’S Disease Neuroimaging Initiative

## Abstract

Although there are many methods available in the literature to compare the covariance structures of two populations, few are suitable for clinical application due to the inability to account for covariate(s) that affect the dependence structure of the variables being investigated. A common method is to adjust the effect of the covariates via a linear model and work with the resulting residuals. However, removing the effects of the covariates could potentially eliminate valuable information from the analysis. We propose a functional nonparametric covariance matrix estimator to account for any given value in the covariate(s), which allows a comparison of the functional covariance structures of the multivariate data. This comparison is facilitated via a test statistic involving the first eigenvalue of the combined form of covariance matrices of the two groups. Three different approaches, namely, the parametric Tracy-Widom, the semi-parametric Forkman’s test, and the nonparametric Permutation method, are used to compute the approximate p-values of the test statistic. We have conducted extensive simulation studies to determine the type I error and power of the proposed hypothesis testing methods and developed practical recommendations for implementing this novel approach. Finally, we apply our methods to the Alzheimer’s Disease Neuroimaging Initiative (ADNI) study to compare cerebrospinal fluid (CSF) biomarkers between dementia and non-dementia cohorts, which offers a fascinating insight into the differences between covariance structures of biomarkers amyloid *β*(1-42) (A*β*42), total tau (tau), and phosphorylated tau (ptau) for given values of age, sex, and years of education.

## 1. Introduction

In clinical settings, a common question is whether a set of predictors significantly influences a binary outcome variable (Thomas, Radji and Benedetti, 2014). Such examples include treatment experiments, gene expressions of phenotypes, and observational studies linking risk factors to a certain outcome, such as dementia. While there is usually only one outcome variable, there could be many, even numerous, variables of interest. In such settings with numerous variables of interest, multiple hypotheses testing methods would inevitably result in a high false positive rate (Storey, 2002). On the other hand, applying corrections to multiple hypotheses testing would inevitably apply severe penalties to statistical power (Blakesley et al., 2009). We propose an alternative solution by creating a single global test that offers greater statistical size and power over multiple testing via comparing the equivalence of the covariance matrices between two groups (Frost, Amos and Moore, 2016). The differences between covariances can provide insight into the dependence structure of the variables being studied, which often can be more informative than a comparison of means. Our approach involves combining the covariance matrices of both groups, creating a test statistic based on its first eigenvalue, and using parametric and nonparametric methods to calculate the p-values. In principal component analysis (PCA), eigenvalues correspond to the amount of variance explained by the components (Wold, Esbensen and Geladi, 1987). By testing the first eigenvalue, we can use a single hypothesis testing framework and bypass the false positive issues stemming from multiple tests.

The current work has two key contributions. First, this paper proposes three hypothesis testing methods for comparing covariance matrices: Tracy-Widom distribution, permutation test, and Forkman’s test. The Tracy-Widom distribution is a parametric probability distribution of the normalized first eigenvalue of a random Hermitian matrix (Tracy and Widom, 1994). The permutation test uses repeated re-sampling without replacement to simulate a null distribution (Pitman, 1937). The Forkman’s test we introduce assumes a standard Gaussian distribution for the first eigenvalue. Our previous work showed that the Tracy-Widom and Forkman’s test methods work well when the variables of interest are standardized to zero mean and unit variance, although the permutation method has yet to be tested. This paper assumes the data has already been standardized, in which case the true covariance matrix of the variables of interest is equivalent to the correlation matrix.

In clinical settings, it is typically not enough to compare the variables of interest by themselves, but it also needs to control for common confounding covariates such as age, sex, body mass index (BMI), etc. (Janes and Pepe, 2008). A confounding covariate is 1) associated with the outcome, 2) associated with the variables of interest, and 3) not be an effect of the variables of interest nor be a factor in the causal pathway of the outcome (Jager et al., 2008). Popular ways of dealing with confounding covariates include randomization, matching, stratification, and restriction (Jager et al., 2008). These approaches aim to remove the confounding effects to assess the effect of the variables of interest on the outcome in a vacuum. However, in the real world, these effects do not happen in a vacuum, and removing these confounding effects could also remove important information. Therefore, the other novel contribution of the current work is to incorporate the covariates into the analysis to observe how given values of the covariates influence the results. To achieve this goal, we use a nonparametric approach to estimate a functional covariance matrix for each given value of the covariates (Yin et al., 2010).

A practical clinical implementation of our methods is to reduce the risk of CSF inflammation, which has been shown to be related to the onset of dementia (Morgan et al., 2019; Pasqualetti, Brooks and Edison, 2015; Hopperton et al., 2018). Both CSF inflammatory biomarkers and dementia progression could be related to confounding covariates such as age, sex, and education level. Traditional methods would remove the covariates’ effects; however, valuable knowledge could be gained from examining inflammation and dementia for different values in the covariates. Our methods would allow for such an examination of continuing covariates without the need to create dichotomized groups. We explore this clinical example by analyzing the Alzheimer’s Disease Neuroimaging Initiative (ADNI) dataset (Jack Jr et al., 2008).

In section 2, we describe in depth the theoretical details of the functional covariance estimation. We provide these details for different covariate situations, i.e., a single continuous covariate, a combination of continuous and categorical covariates, and multiple continuous covariates. Section 3 explains the theoretical backings and implementation of the aforementioned covariance comparison methods. The nuances and drawbacks of each method based on previous works are also discussed in this section. Section **??** details the simulation designs for both single and multiple covariate(s) simulations, followed by an evaluation of type I error and power for repeat simulations of said designs. In section 4 we apply our methods to the ADNI dataset. Our objective in the data analysis is to determine whether there are differences in CSF biomarkers between dementia and non-dementia participants after adjusting for significant covariates. Finally, section 5 discusses our results, limitations, and future directions for this work.

## 2. Nonparametric conditional covariance function estimation

The conditional covariance matrix for a given value(s) in the covariate space can be estimated via a nonparametric functional estimator (Yin et al., 2010). For two random vectors **x** and **u**, the conditional covariance of **x** given a fixed value of **u** is represented as Cov(**x** | **u**) = **Σ**(**u**). Where **x** = (*x*_1_, …, *x*_*p*_)^*′*^ ∈ ℝ^*p*^ and **u** = (*u*_1_, …, *u*_*q*_)^*′*^ ∈ ℝ^*q*^. The main focus of the paper will be on *q* = 1, with a section exploring the inclusion of a categorical covariate and another section exploring the *q* = 2 case. High dimensional *q* runs into the problem of the curse of dimensionality in nonparametric estimation that requires additional workarounds (Lavergne and Patilea, 2008), an approach will be briefly discussed in the Discussion section.

### 2.1. One-dimensional covariate

Given vector **x**_*i*_ = (*X*_*i*1_, …, *X*_*ip*_)^*′*^, and scalar *U*_*i*_, *i* = 1, …, *n* be random samples from (**x**, *U*). Let *K*(*u*) be a symmetric kernel density, and *K*_*h*_(*u*) = *h*^*−*1^*K*(*u/h*) where *h* is a positive bandwidth (Yin et al., 2010). The local estimators of conditional mean (**m**(·)) and covariance (**Σ**(·)) are the minimizers of the objective function

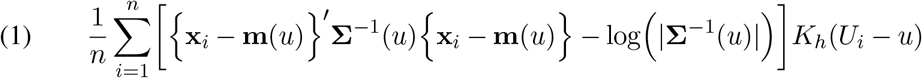

The minimizers of (1) are the estimators:

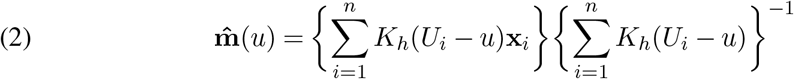

and

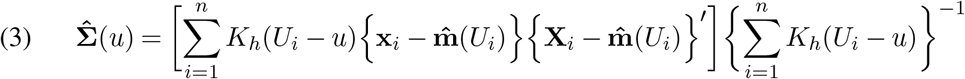

The mean estimator 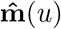 is widely known as the local constant estimator. In practice, we prefer the local linear estimator as it is robust to bias, especially near the boundaries of the estimation range (Cheruiyot, 2020).

Briefly, the local linear estimator for the mean is given by:

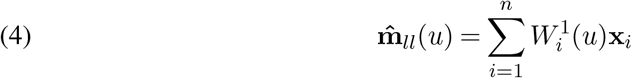

where

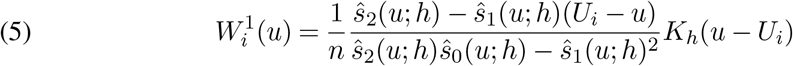

and

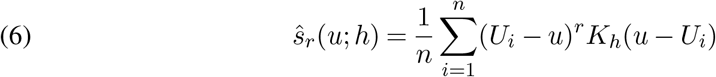

However, the local linear estimator could not be directly used for 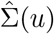 as it could produce invalid covariance matrices that are not positive semi-definite without additional workarounds. A local linear estimator method facilitated by the modified Cholesky decomposition (MCD; Chen and Leng, 2015) will be discussed in the Discussion section.

A method to select the optimal bandwidth for 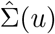 is by minimizing the log likelihood leave-one-out criterion:

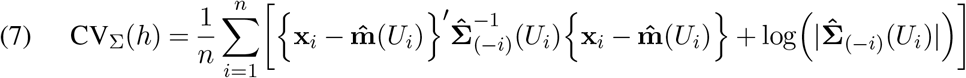

Where 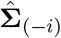 is the covariance estimator computed without the *i*th data point. The optimizer function is computationally expensive if the sample size is large, as the leave-one-out crossvalidation method calculates the criterion once for each sample. The covariance estimator can estimate the conditional covariance matrix for any given *u* ∈ *U*, but to avoid extrapolation, we highly recommend estimating only within the range of the available data.

A correlation matrix can be defined as a special type of covariance matrix where the variables have been standardized to unit variance. However, the nonparametric conditional covariance estimator cannot be viewed as a true correlation estimator, as it does not force the diagonals to equal 1. While the methods in this paper require standardization of the data, we will nevertheless refer to both the estimator and true covariance/correlation matrix as a covariance matrix for consistency unless the distinction is necessary.

### 2.2 One-dimensional covariate with a categorical covariate

This section will develop an extension of the conditional covariance estimator, incorporating a categorical and continuous covariate. Often, categorical covariates are important factors to adjust for in clinical settings, a popular example being the gender of a subject.

The local linear regression estimator of the mean can incorporate categorical variables by adapting the Aitchison and Aitken unordered discrete kernel (Aitchison and Aitken, 1976), where a higher weight is given to samples having the desired factor level. The Aitchison and Aitken kernel can also be applied directly to conditional covariance matrix estimation.

Let *U*_1_ be a continuous variable, *U*_2_ be a categorical variable, and *c* be a given factor level of *U*_2_, the covariance estimator is given by:

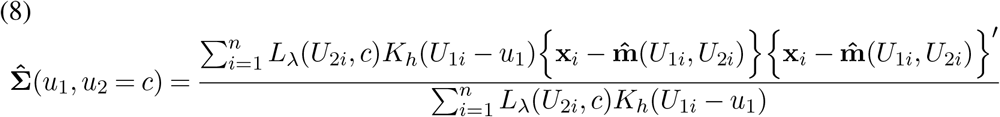

Where

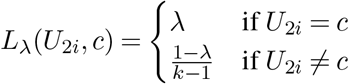

*k* is the number of factor levels, and *λ* ∈ [1*/k*, 1] is the tuning parameter indicating the weight given to the desired factor level. In the data analysis, we set *λ* = 1.

While the local mean estimator incorporates the entire sample, the covariance estimator only uses samples that match the factor level of the categorical covariate at which the covariance is estimated. Therefore, ensuring that each factor level has enough data points for an accurate estimation is necessary for the proposed approach.

### 2.3. Multi-dimensional covariates

Extending the conditional covariance estimator to a *q* dimensional (**U** ∈ ℝ^*n×q*^) is straightforward. The local linear mean estimator can be extended from the one-dimensional case to multiple dimensions by replacing equation (5) with a vectorized version (García-Portugués, 2024):

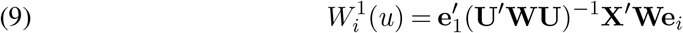

where *W* = diag(*K*_*h*_(*U*_1_ − *u*), …, *K*_*h*_(*U*_*n*_ − *u*)), 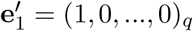, and **e**_*i*_ is a column vector with 1 as the *i*th entry and 0 otherwise.

The covariance estimator can be extended to multivariate situations by using the product kernel, also known as the Nadaraya-Watson (NW) kernel estimator (Cai, 2001):

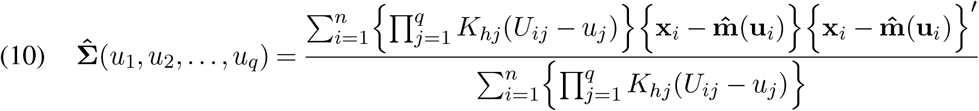

Both the mean estimator and covariance estimator would suffer from the curse of dimensionality, where there are few data points within a reasonable bandwidth unless there is an overabundance of data. A possible solution for the mean estimation is to use the generalized additive model (GAM; Hastie, 2017), an approach discussed in the conclusions chapter. But for small dimensions such as *q* = 2, the curse of dimensionality is unlikely to be an issue given an adequate sample size.

## 3. Two-group covariance comparisons

In the previous section, we have introduced a nonparametric estimator of the conditional covariance function. In this section, we will develop a range of tools to compare the covariance matrices of two groups. For a fixed covariate value at *u*, we will discuss hypothesis testing methods to test

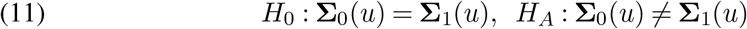

Where **Σ**_0_(*u*) and **Σ**_1_(*u*) are the covariance matrices of **X**_0_ and **X**_1_ conditioned on **u**.

To test the hypothesis, we will introduce the Tracy-Widom, Forkman’s test, and permutation methods in the following subsections. We standardize the data in order to use the aforementioned methods for testing correlation matrices.

### 3.1. Tracy-Widom Approach

The Tracy-Widom distribution is the probability distribution of normalized eigenvalues from a random Hermitian matrix (Tracy and Widom, 1994). The Tracy-Widom distribution (of order 1) has many applications in multivariable analysis; of relevance is the two-group comparison of covariance matrices (Johnstone, 2008, 2009). In our work, we utilize the Tracy-Widom hypothesis method to compare conditional covariance across two groups in the following way:

Given two sample groups **D**_0_ ⊂ (**X**_0_, **U**_0_) and **D**_1_ ⊂ (**X**_1_, **U**_1_), let 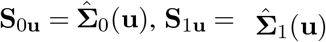. Furthermore, (*n*_0_, *n*_1_) are the sample sizes of (**D**_0_, **D**_1_) respectively, and *p* the number of variables of interest.

Then

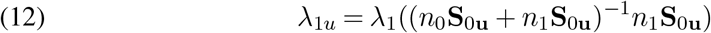

where *λ*_1_(·) is the first eigenvalue of a given positive semi-definite matrix.

Therefore, the test statistic, which follows the Tracy-Widom distribution of order 1, is

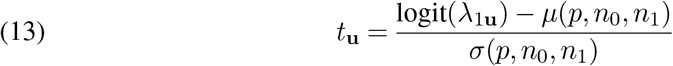

*μ* and *σ* are, respectively, the centering and scaling terms defined as:

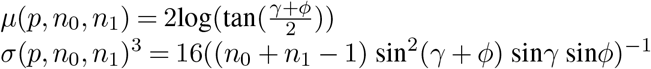

where

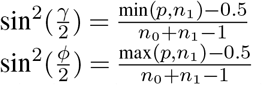

The hypothesis testing has a one-sided upper-tail p-value given by *Pr*(*T > t* | *H*_0_) = 1 − *F*_1_(*t*), where *F*_1_(*t*) is the cumulative distribution function of the Tracy-Widom distribution under *H*_0_. As the actual Tracy-Widom p-value is challenging to calculate directly, a Gamma distribution approximation is instead used to calculate the p-value (Chiani, 2014).

For the Tracy-Widom distribution to be valid, various regularity conditions must be met, such as both groups are independent and follow Gaussian distribution. Furthermore, the Tracy-Widom distribution is an asymptotic distribution (Frost, Amos and Moore, 2016), defined as 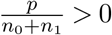 and 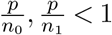 as *p, n*_0_, *n*_1_, → ∞. However, simulations have shown that Tracy-Widom serves as a reasonable if not excellent approximation for small and moderate *n*_1_, *n*_2_, and *p* (Johnstone, 2001). In addition, further theoretical results reveal that Tracy-Widom holds true for non-Gaussian data under a set of conditions (Soshnikov, 2002). These conditions include the data having zero mean and unit variance, a symmetric distribution, and the tails of said distribution approaching zero as fast as the Gaussian distribution.

In practice, the zero mean and unit variance conditions are satisfied by data standardization, and transformation can be used to coerce the data to be roughly symmetric.

### 3.2. Forkman’s test

In this section, we introduce a novel hypothesis testing method for calculating p-value by generating the bootstrap samples from standard Gaussian distributions, coined the Forkman’s test. The original Forkman’s test is a bootstrap algorithm that determines the number of significant principal components in a standardized data matrix (Forkman, Josse and Piepho, 2019). We take inspiration from their work, particularly the generation of bootstrap samples from standard Gaussian distributions. We use the first principal component to determine and identify differences in covariance matrices via bootstrap. Our modified Forkman’s test algorithm is as follows:

Given *λ*_1**u**_ as defined in Section 3.1, the test statistic is

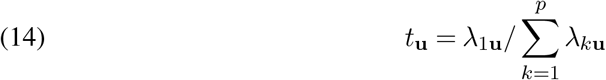

Where **Z**_**u**_ = (*n*_0_**S**_0**u**_ + *n*_1_**S**_1**u**_)^*−*1^*n*_1_**S**_1**u**_ and *λ*_*k***u**_ = *λ*_*k*_(**Z**_**u**_).

The Forkman’s test p-value is calculated using the following algorithm:

### 3.3. Permutation test

The permutation test is a resampling method that randomly shuffles the outcome group labels without replacement to generate a new set of observations (Ojala and Garriga, 2010). To avoid introducing new bias from estimating the mean function for each resampled data, we estimate the means on the original data before shuffling the group labels. Afterward, the estimated residuals are shuffled, and the bootstrap covariance matrices are calculated using the shuffled residual. We combine the covariance matrices to find the bootstrap test statistic for each iteration; these test statistics are then aggregated to calculate the bootstrap p-value.

#### Algorithm 1

Forkman’s Test P-Value

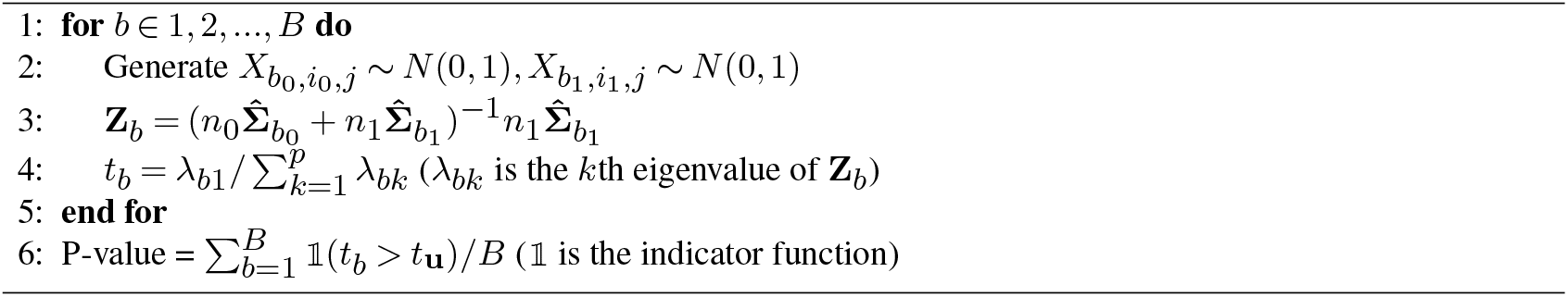

#### Algorithm 2

Permutation P-Value

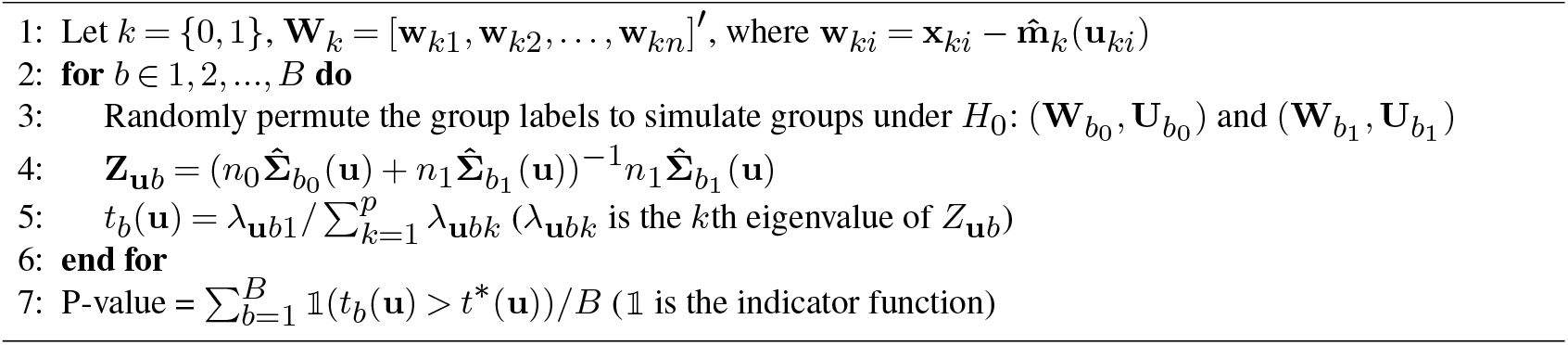

Note that while theoretically, the optimal bandwidths for the covariance matrices should be recalculated for each bootstrap data, in practice, it is too computationally expensive to be feasible. Therefore, for practicality, we use the same bandwidths as the original covariance matrices in each bootstrap iteration.

### 3.4. Simulation design

In this section, we have performed extensive simulations to study the performances of the proposed methods. We designed simulation studies for univariate and bivariate continuous covariate cases.

### 3.5. One-dimensional continuous covariate

We use a simple simulation of two predictors *X* = (*X*_1_, *X*_2_)^*′*^ and one covariate *U* with equal sample size *n* for two groups *j* = 1, 2.

Simulate

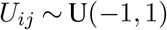

Let

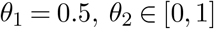

Simulate

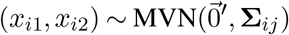

where

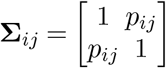

and

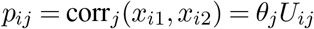

### 3.6. Two-dimensional continuous covariate

We make the following modification to the simulation design in section 3.5 for two-dimensional continuous covariates *U* :

Simulate

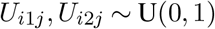

Let

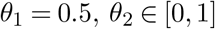

Simulate

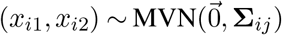

where

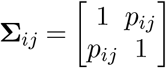

and

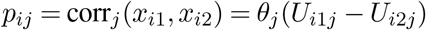

We use the simulation design to calculate the type I error and power of the three hypothesis testing methods’ introduced in the last section. We set both groups to have equal sample sizes (*n*_1_ = *n*_2_ = 500) and 100 repetitions. Type I error is calculated by setting *θ*_1_ = *θ*_2_ = 0.5, while power is plotted for *θ*_2_ = 0 and *θ*_2_ = 1. In theory, the covariance matrices should be the same when *U*_1_ = *U*_2_, and thus, the power plots should represent type I error for these cases.

### 3.7. Simulation Implementation and Results

Section 3.7.1 - 3.7.3 provides the results for the one-dimensional simulation and section 3.7.4 for the two-dimensional simulation.

#### 3.7.1. First eigenvalue accuracy

To assess the accuracy of the first eigenvalue, which is used for the test statistic, of the estimated covariance matrix, we compare the actual curve of the first eigenvalue (*λ*_1u_) to the estimated first eigenvalue 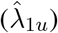 of the combined matrix.

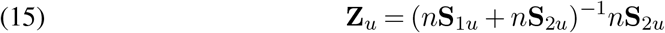

We have performed this simulation using 50 repetitions and *n* = 5000, and cross-section plots are presented for *θ* ∈ (0, 0.5, 1). The bandwidth used is the minimizer of the log-likelihood leave-one-out criterion (equation 7).

One observation of Figure 1 is that the estimated first eigenvalue is smoother and seemingly more accurate to the true first eigenvalue when *θ*_2_ is small, and the opposite is true when *θ*_2_ is large. A possible explanation for a small *θ*_2_ leading to smoother estimates is that the corresponding simulated covariance matrices are close to the identity matrix for small covariance values. In addition, the estimated function seems closer to the true eigenvalue function when *θ*_2_ is small; perhaps the noise associated with estimating higher correlated covariance matrices negatively impacts the estimates. Also, when *θ*_1_ = *θ*_2_ = 0.5, the true eigenvalue of the combined matrix is not dependent on *U*, but is a flat value of 0.5. While the estimated eigenvalue is essentially flat, it is also marginally higher for all values of *U*. Lastly, it is interesting that small values of *θ*_2_ underestimate the eigenvalue near the edge while overestimating near *U* = 0. When *θ*_2_ is large (0.5 and higher), the estimates overestimated the eigenvalues for all values of *U*. Overall, the study shows that the estimates of the first eigenvalue provide a good approximation to the true value as a function of *U*.

**FIG 1.**
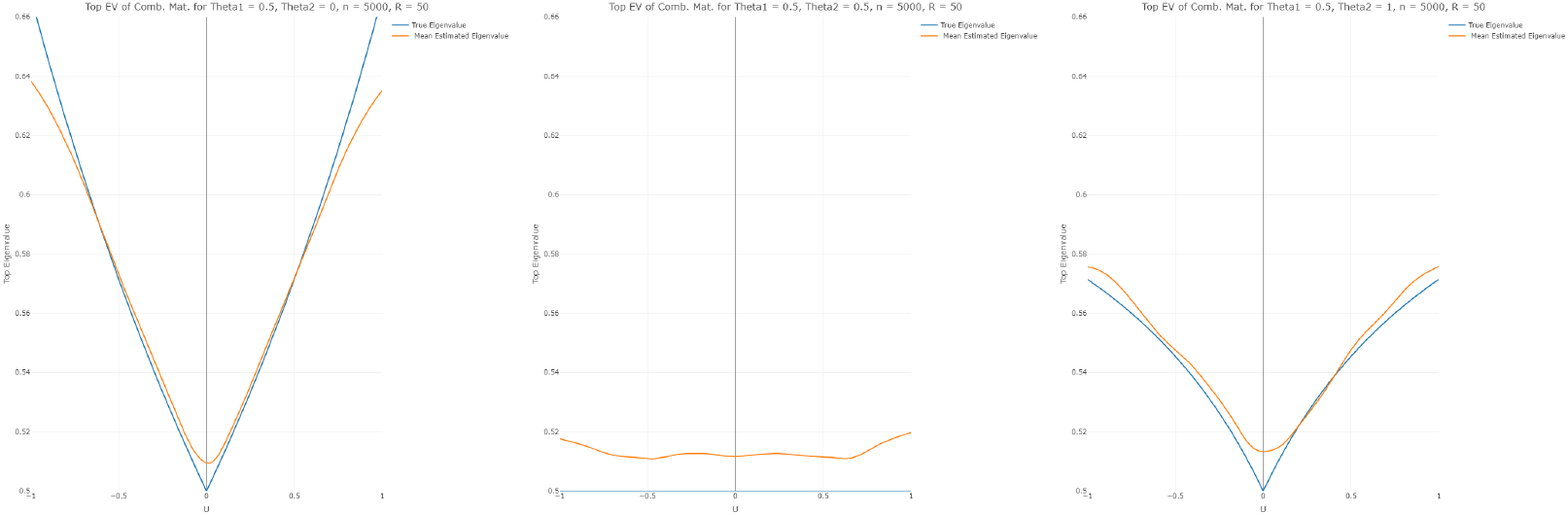
True vs estimated first eigenvalue of Z_u_

**FIG 2.**
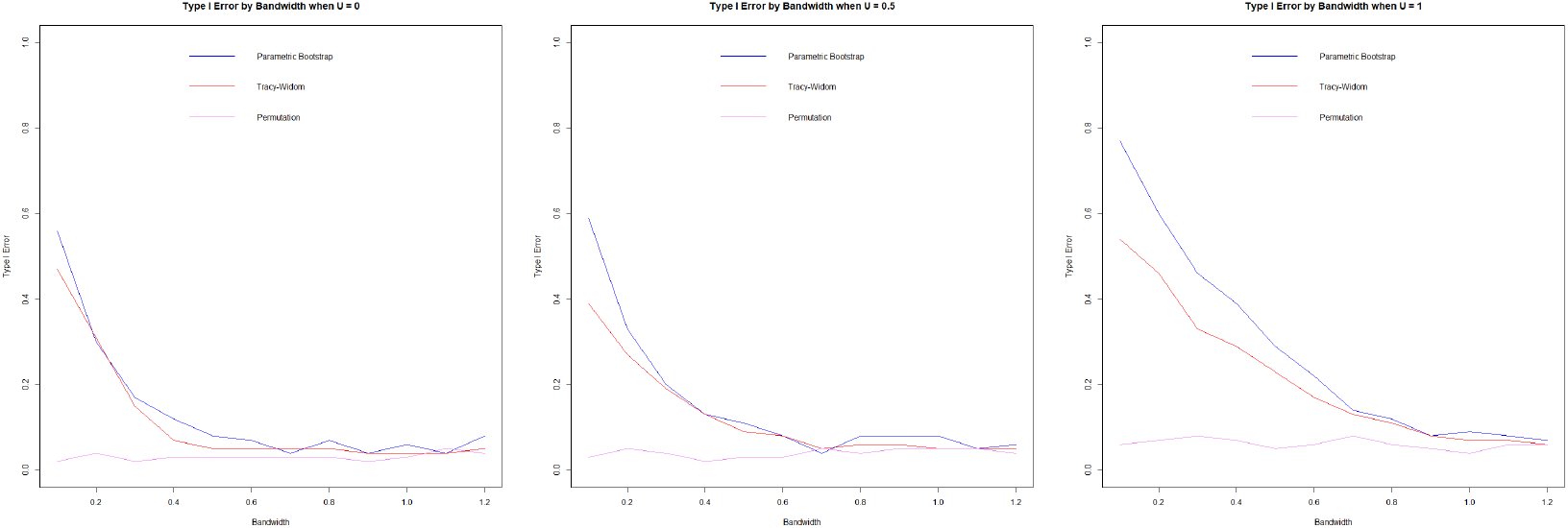
Type I error by bandwidth (h), U = 0 (left), U = 0.5 (center), U = 1 (right)

**FIG 3.**
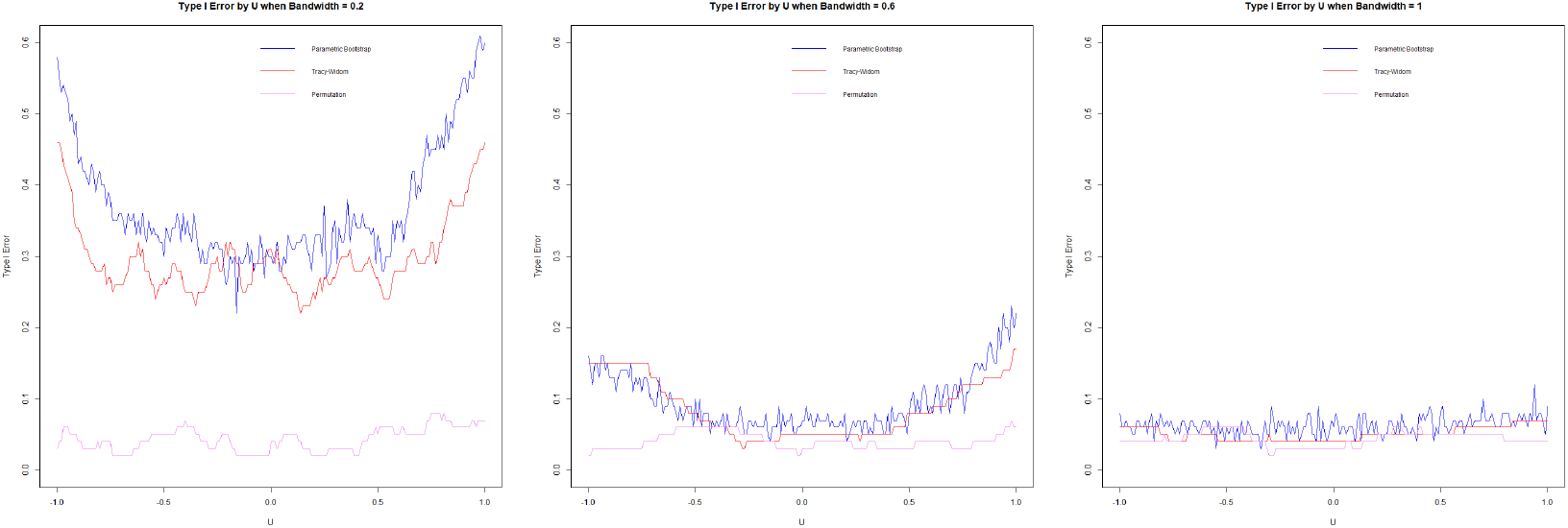
Type I error by U, h = 0.2 (left), h = 0.6 (center), h = 1 (right)

**FIG 4.**
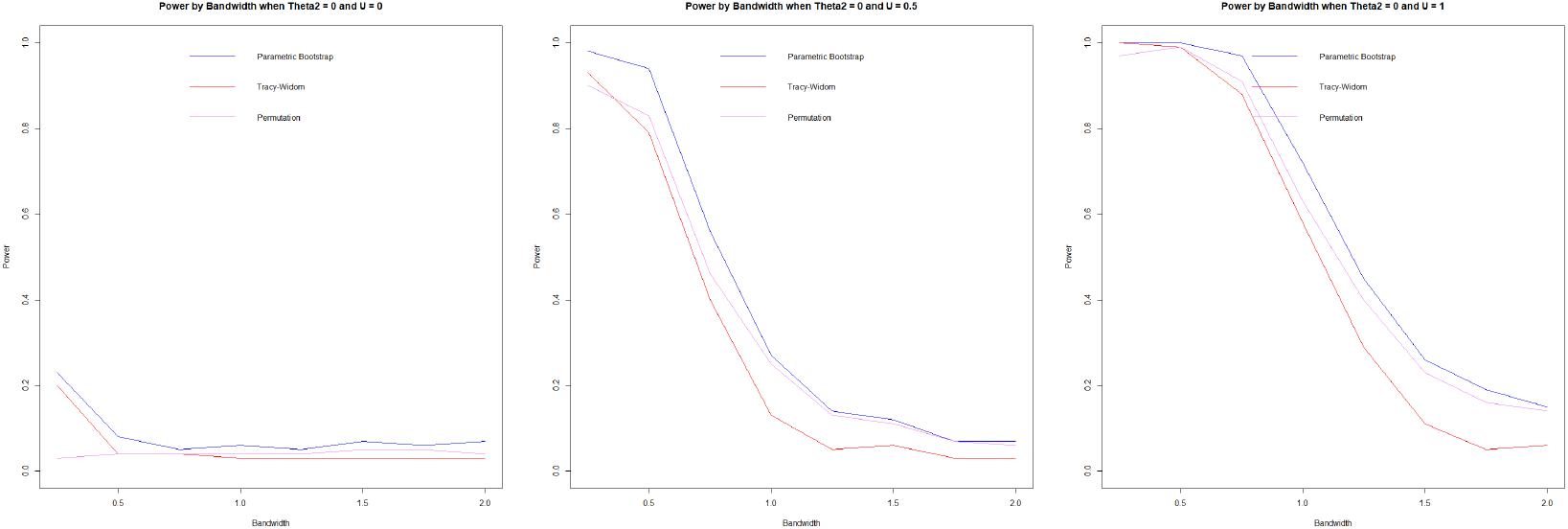
Power by bandwidth, θ_2_ = 0, U = 0 (left), U = 0.5 (center), U = 1 (right)

**FIG 5.**
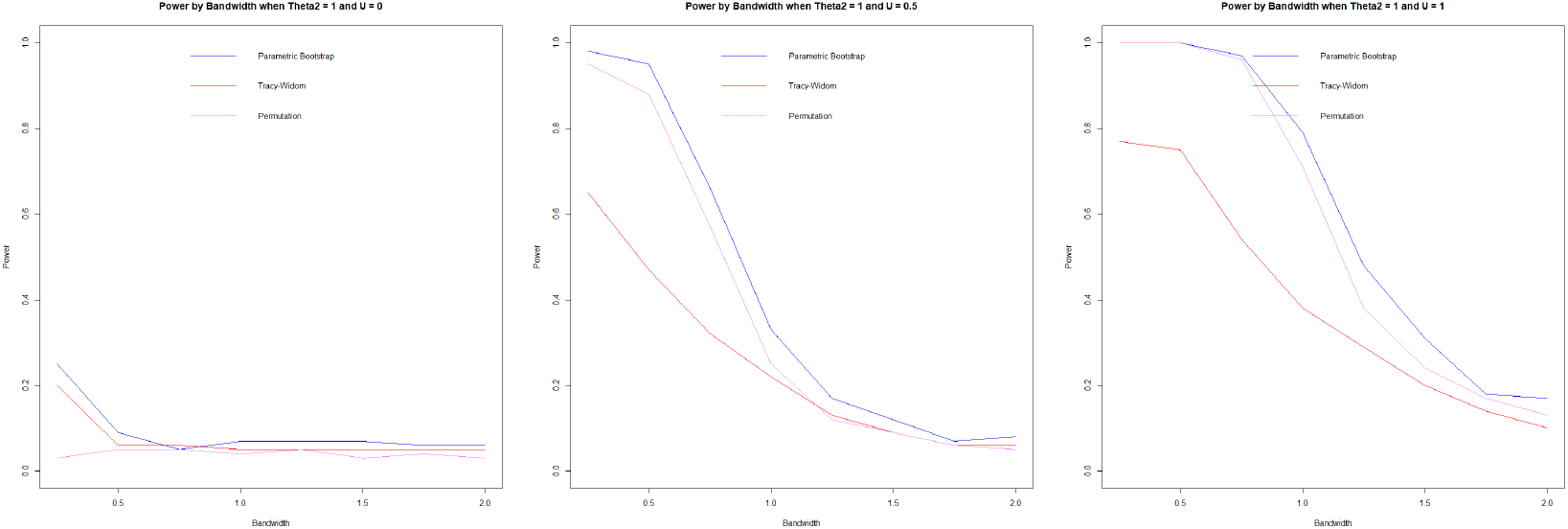
Power by bandwidth, θ_2_ = 1, U = 0 (left), U = 0.5 (center), U = 1 (right)

**FIG 6.**
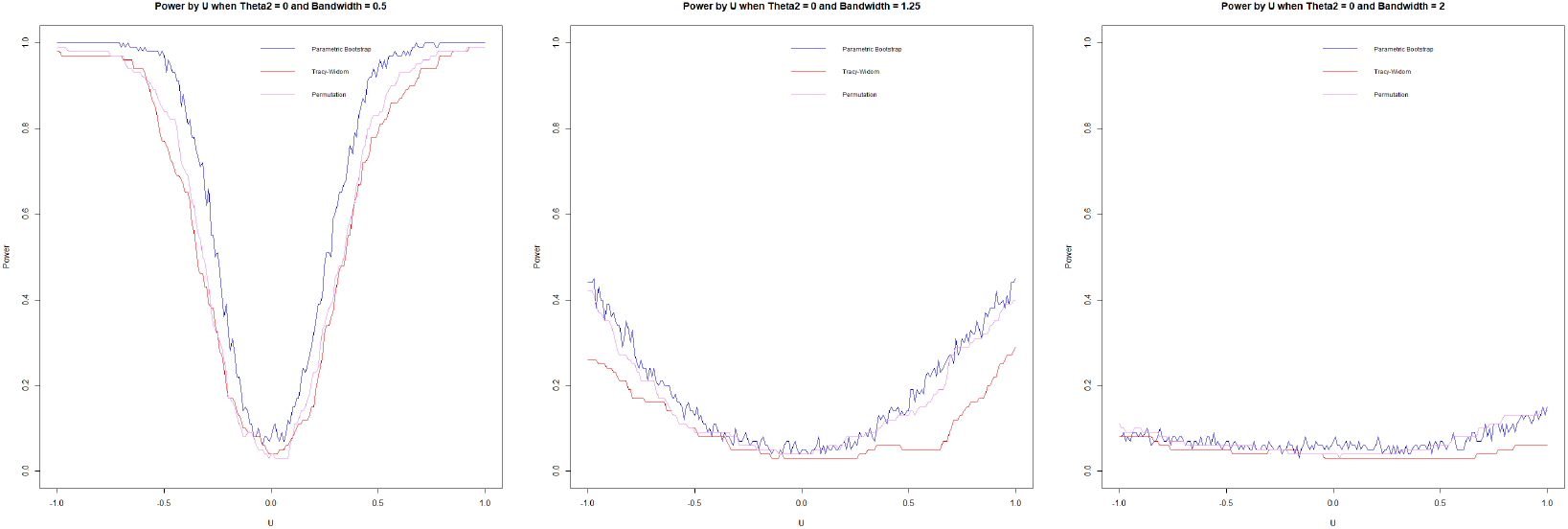
Power by bandwidth, θ_2_ = 0, h = 0.5 (left), h = 1.25 (center), h = 2 (right)

**FIG 7.**
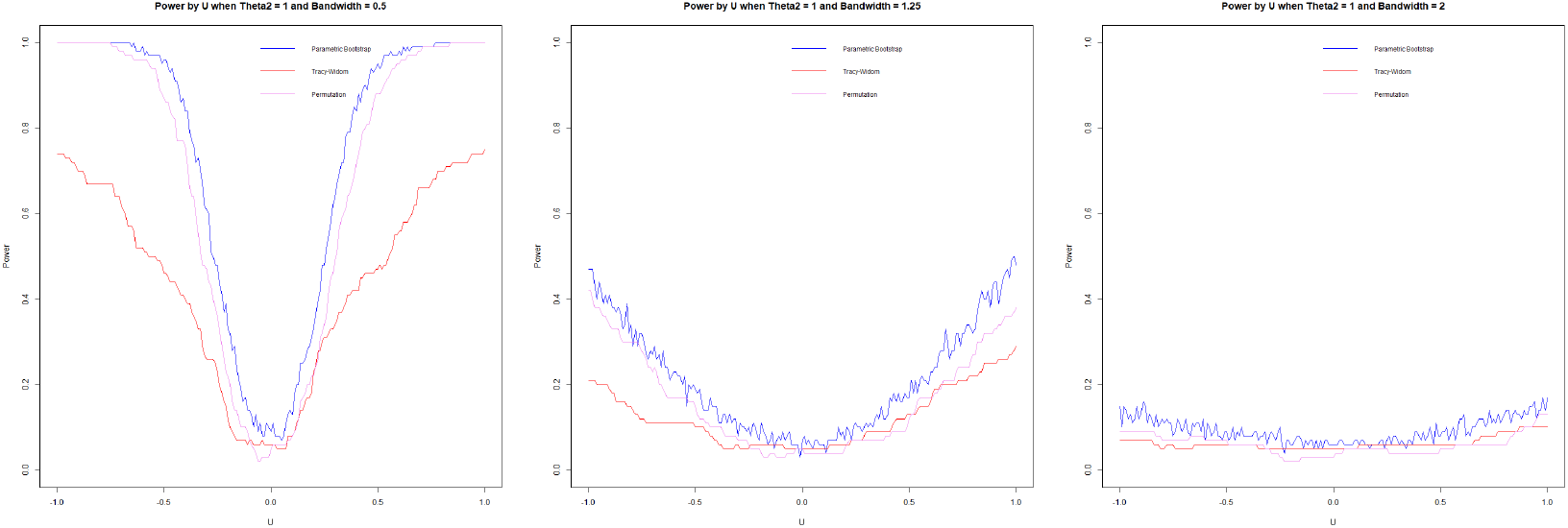
Power by bandwidth, θ_2_ = 1, h = 0.5 (left), h = 1.25 (center), h = 2 (right)

#### 3.7.2. Type I error

We assess the type I error by simulating from models in Section 3.5 with *θ*_1_ = *θ*_2_ = 0.5, using 100 repetitions and *n* = 1000. We visualize the Type I error via significance trace plots by plotting Type I error curves as a function of a range of bandwidths (*h*) and *U*.

The significance trace plots show that the permutation method is extremely robust, accurately estimating Type I error for all values in *U* and the bandwidth (*h*). But Tracy-Widom and Forkman’s test methods struggle with high Type I error when *h* is small. These methods are based on parametric distributions (Tracy-Widom on Wishart and Forkman’s test on Gaussian); as such, they could struggle with high variance for small *h*.

#### 3.7.3. Power

Power is assessed for *θ*_2_ = 0 and *θ*_2_ = 1 using 100 repetitions and *n* = 1000. When *u* = 0, the covariance matrices of both groups are equal regardless of the values of *θ*_2_; as such, it represents type I error when *u* = 0.

While a higher *h* is advantageous for minimizing type I error, a lower *h* is advantageous for maximizing power. Forkman’s test and permutation methods have very similar power results, with Forkman’s test having slightly higher power and type I error. In our simulation design, the Tracy-Widom test underperformed, having comparatively lower power while not improving the type I error. Permutation and Forkman’s test are strong performers for the power simulation.

In the null case, the bandwidth, as the cross-validation suggests, is smaller than the *h* that accurately attains the targeted type I error. In the alternative case of *θ*_2_ = 0, the optimal *h* (very high) would lead to low power, and a smaller *h* would improve power. The opposite is true for *θ*_2_ = 1, where the optimal *h* is too low. While the test is powerful, it causes a high type I error that could be mediated with a higher *h*. In summary, cross-validation (equation 7) does not select optimal *h* in these cases, which suggests that the problem calls out for other bandwidth selection methods that we have not investigated in this work. We suggest some potential improvements in the discussion section (Section 5).

#### 3.7.4. Multiple covariates results

The simulations clearly show that Forkman’s test is the most powerful, followed by the Tracy-Widom and permutation methods (Figure 8). However, in these simulations, the high power comes with the downside of high type-I error. Therefore, the Forkman’s test method is good when the priority is to maximize power, the permutation method is suitable for minimizing type I error, and the Tracy-Widom strikes a balance between type I error and power. An interesting observation is that when *θ*_2_ = 0, the Tracy-Widom and Forkman’s test has good power while the permutation method does not. This result contrasts with the single covariate results where the permutation method closely resembles Forkman’s test method and outperforms the Tracy-Widom method in power. When *θ*_2_ = 1, Tracy-Widom and Forkman’s test methods have very high type I error, while only the permutation method has a low type I error. The permutation test struggles in multidimensional scenarios as it is a nonparametric method free of rigid parametric assumptions but suffers from inefficiency (Ramdas et al., 2014). In summary, each method can be the best performer given the circumstances; there is no one method that is better or worse than others.

**FIG 8.**
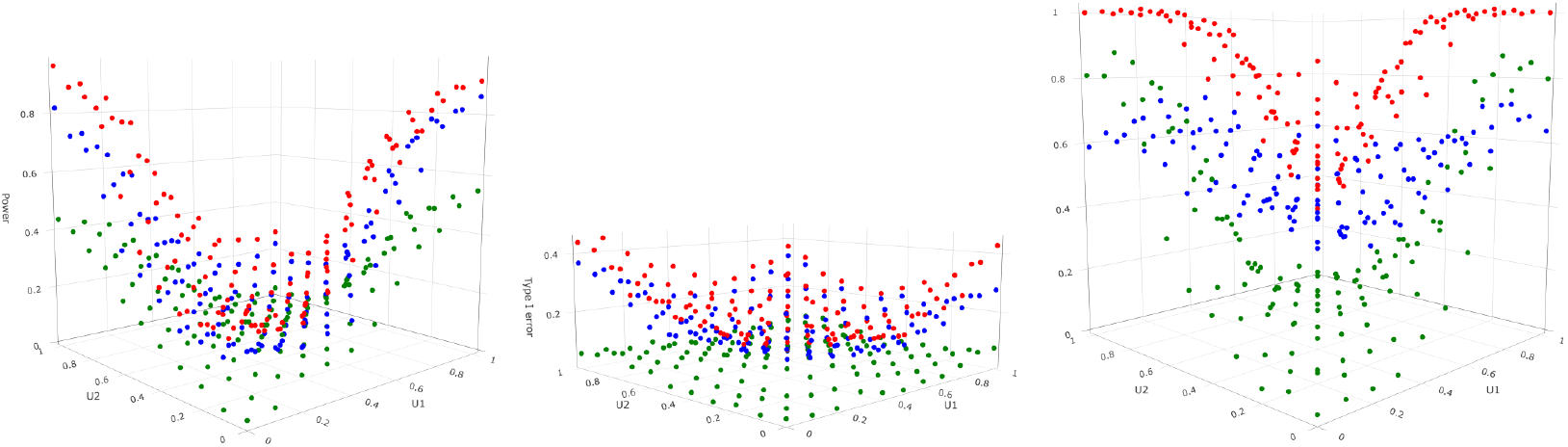
2D Covariates Simulation: Power by U_1_ and U_2_; θ_2_ = 0 (left), θ_2_ = 0.5 (type I error; center), θ_2_ = 1 (right). Power and type I error are calculated for Tracy-Widom (blue), Forkman’s test (red), and permutation (green)

## 4. ADNI biomarkers and dementia data analysis

### 4.1. Methods

To assess the practicality and usage of conditional covariance comparison, we analyzed data from the Alzheimer’s Disease Neuroimaging Initiative (ADNI) database (adni.loni.usc.edu). The ADNI was launched in 2003 as a public-private partnership led by Principal Investigator Michael W. Weiner, MD. The primary goal of ADNI has been to test whether serial magnetic resonance imaging (MRI), positron emission tomography (PET), other biological markers, and clinical and neuropsychological assessment can be combined to measure the progression of mild cognitive impairment (MCI) and early Alzheimer’s disease (AD). For up-to-date information, see www.adni-info.org. The ADNI database provides access to wide-ranging clinical, imaging, genetic, and biomarker data (Mueller et al., 2005). Among the biomarkers data are CSF biomarkers amyloid *β*(1-42) (A*β*42), total tau (tau), and phosphorylated tau (ptau). Studies have shown that A*β*42 is a strong predictor of dementia progression (van Harten et al., 2013), that tau can be an aid in AD diagnosis (Mulder et al., 2010), and ptau has excellent retrospective diagnostic performance for AD and other forms of dementia (Thijssen et al., 2021). While these results are meaningful, they did not consider the important risk factor of age. We aim to condition the CSF biomarkers values on age at the time of testing and evaluate their significance in predicting lifetime dementia progression.

The baseline diagnosis was taken from the first examination for each participant, regardless of which ADNI study the participant first enrolled in. Meanwhile, the follow-up dementia diagnosis is not observed at a fixed time interval but as the last available diagnosis in the study. A positive diagnosis of dementia was considered a positive outcome, while a negative diagnosis and mild cognitive impairment (MCI) are considered negative outcomes. As we are interested in observing lifetime dementia progression, participants with positive outcomes at baseline were dropped (n = 229). Additionally, we removed participants whose last diagnosis was within three years of baseline diagnosis (n = 175) to improve the quality of follow-up diagnosis. Extreme biomarker values were not numerically measured and thus were truncated to a fixed value. The CSF biomarkers distributions exhibit extreme right-skewness, violating the Gaussian distribution assumption of the covariance estimation. Thus, we use the common practice of log transformation to improve symmetry and subsequently standardize the values (Changyong et al., 2014). Conditional covariance matrix functions were calculated for both diagnosis outcomes, using optimal bandwidth via cross-validation based on the loglikelihood leave-one-out criterion. Finally, test statistics and p-value at (*α* = .05) significance level are calculated for a set age interval (55 years to 90 years by 0.5 years) using the three aforementioned methods.

### 4.2. Biomarkers by age

The ADNI analytic data has (n = 811) records, with (n = 590) having negative follow-up outcomes and (n = 221) positive dementia outcomes. The optimal bandwidth for covariance estimation was 8 years for negative outcomes and 6 years for positive outcomes. The results show that the covariance matrices for the two groups generally differ for ages 77 and under (Figure 9). Specifically, the Forkman’s test method was significant for ages under 80.5 and above 85, and Tracy-Widom was significant for under 75 and over 89.5. In contrast, the permutation method was significant for ages between 59 and 79.5 and 88.5 and above. The p-value trend is highest for Tracy-Widom, followed by permutation, with Forkman’s test having the lowest.

**FIG 9.**
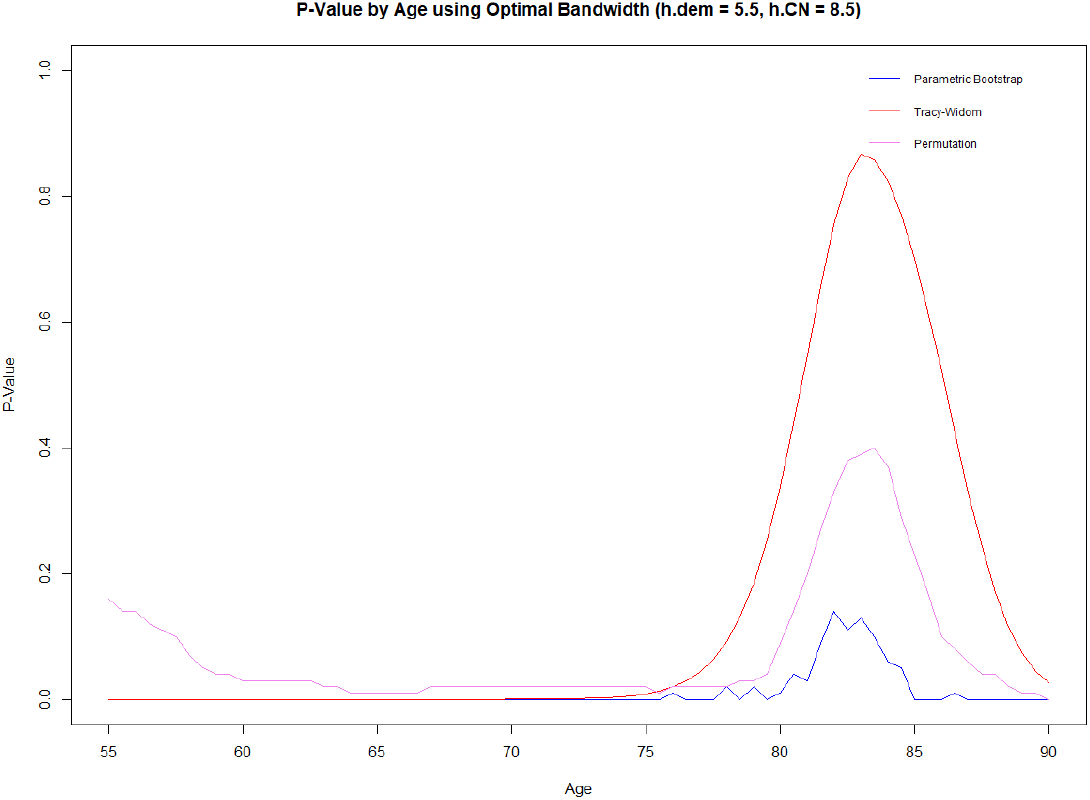
ADNI data analysis: Comparison of p-values by age

**FIG 10.**
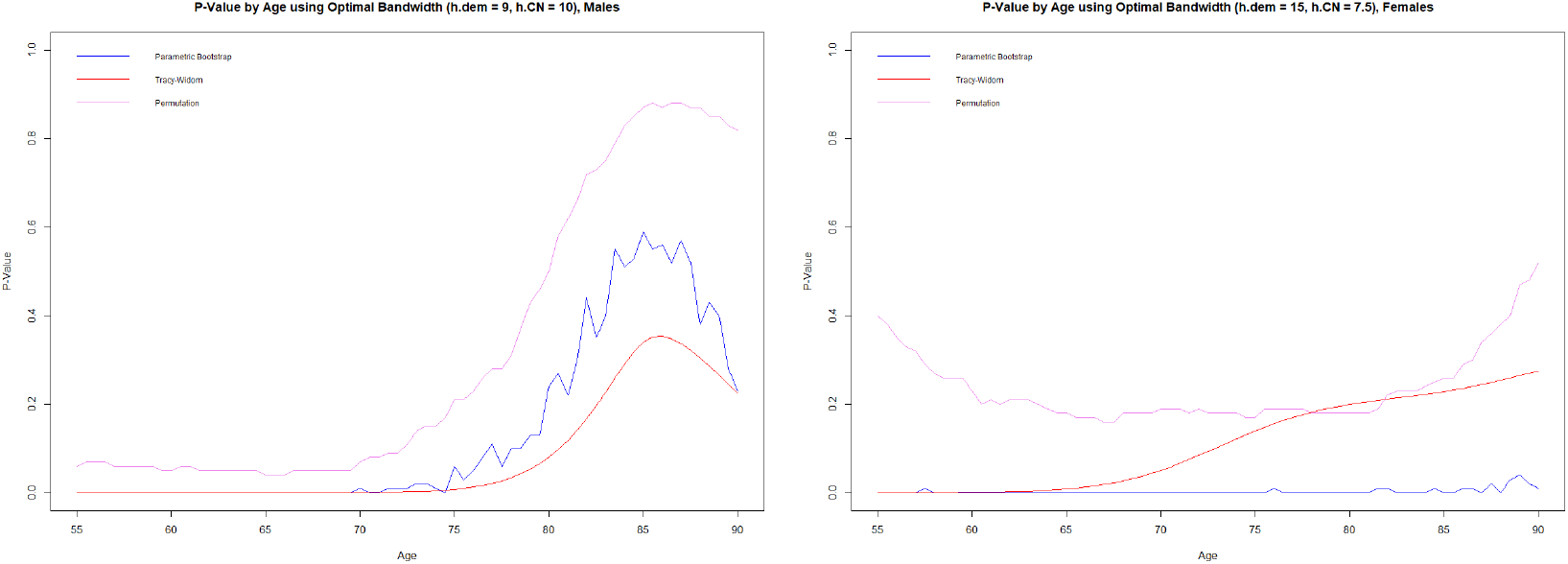
P-values for males (left) and females (right)

The data analysis results show differences in covariance matrices between the dementia and non-dementia groups for certain age groups. Given that nonparametric estimation at the borders of the age range can often be biased (Hall and Wehrly, 1991), we can not adequately assess the results near the ages of 55 and 90. However, we can confidently conclude that the covariance matrices of CSF biomarkers significantly differ between the ages of 59 and 77. These results suggest that the covariance matrices of CSF predictors A*β*42, tau, and ptau can together be useful in predicting dementia progression for younger subjects. This result gives even more importance to the necessity for early detection of dementia and AD when the CSF biomarkers can exhibit significant differences. It is important to note that this study examined the interaction effects of the biomarkers but not the main effect. Our work should not be considered as a replacement for established main effect models but rather as a supplement.

### 4.3. Biomarkers by age and sex

To investigate the effect of a person’s sex and age on their biomarkers covariance matrix, we use age as a continuous covariate and sex as a categorical covariate in the method described in Section 2.2. There are slightly more males in the study than females and significantly more non-dementia participants than dementia ones (Table 1). The age’s optimal covariance bandwidth used for males is 9 years for the dementia group and 10 years for the non-dementia group. For females, it is 15 years for the dementia group and 7.5 years for the non-dementia group.

**TABLE 1.** Cross tablulation of sex and dementia outcome.

| Sex\Outcome | Dementia | Non-dementia | Total |
| --- | --- | --- | --- |
| Female | 88 | 281 | 369 |
| Male | 133 | 309 | 442 |
| Total | 221 | 590 | 811 |

The p-value results for males are similar to the overall results, where the p-values are significant for participants below 70 - 80, depending on the method, and not significant otherwise. However, the results for females are much more different, exhibiting a much flatter p-value curve than for the males group. The Forkman’s test p-value is significant for all ages, the Tracy-Widom is significant only for participants younger than 70, while the permutation method is not significant for any age.

While the different methods do not agree on the significance of the p-values in the female cohort, their p-value graphs are flatter overall. This is conclusive evidence that the biomarkers’ conditional covariance matrices for females are much less dependent on age than for males. Possibly this is a result of a smaller sample size for females than males, or perhaps there exists a clinical difference in how the biomarkers (A*β*42, tau, and ptau) change with age between the genders. Indeed, recent works have suggested significant differences between the CSF biomarkers and their interaction with the cognitive outcome of the genders (Mofrad et al., 2020).

### 4.4. Biomarkers by age and education

We investigate the effects of two continuous covariates (age and years of education) via multi-covariate conditional covariance estimation. The distribution of years of education is comparable between dementia and non-dementia groups, both centered at 16 years with a slight left skew (Figure 11). The local linear method was used to estimate the mean and the NW estimator for the covariance, as detailed in section 2.3.

**FIG 11.**
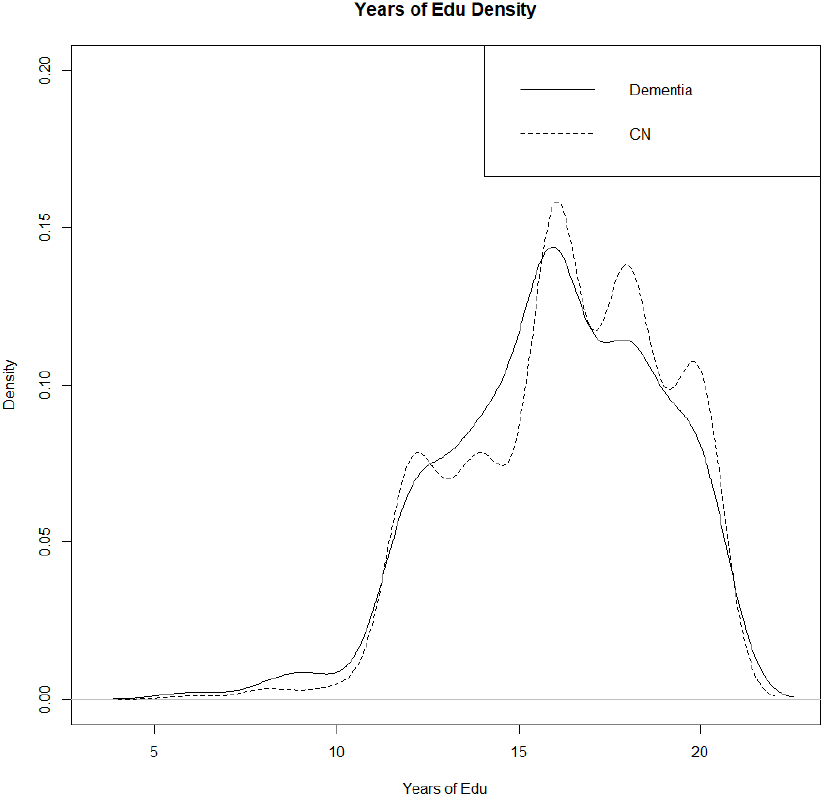
Years of education density

The p-value plots are difficult to visualize due to their dependence on two variables. Our solution uses a top-down view, using red for significant p-values and green for non-significant p-values (Figure 12). The Tracy-Widom and Forkman’s test have similar results, showing significant p-values throughout, aside from ages greater than 75 years and education of above 13 years. However, the permutation p-value is vastly different, only being significant for the age range of 70 - 80 years and not dependent on years of education.

**FIG 12.**
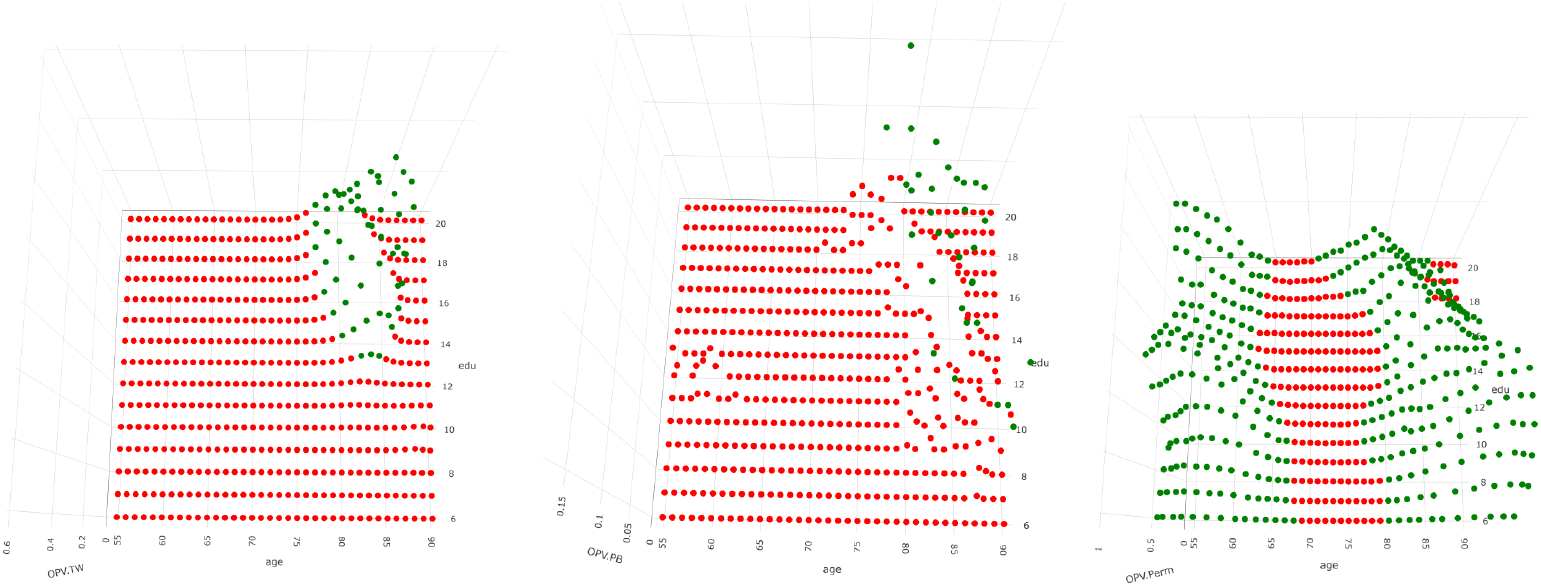
Two-covariate p-values: Tracy-Widom (left), Forkman’s test (center), Permutation (right)

The results suggest that the Tracy-Widom and Forkman’s test methods are powerful in finding significant covariance matrices, while the permutation method is less powerful in comparison. The Tracy-Widom and Forkman’s test results show that both age and years of education are important factors in identifying covariance differences in CSF biomarkers, while the permutation method suggests only age is necessary.

## 5. Discussion

### 5.1. Future Work

There is potential room for improvement in the conditional covariance matrix estimation and comparison research. For one, the local linear estimator outperforms the local constant estimator with respect to bias in the boundaries of the kernel and asymptotic mean squared error at the cost of higher computational needs (Yu and Jones, 1997). However, directly applying the local linear estimation to the conditional covariance matrix would not guarantee the positive semi-definiteness of the estimator. Instead, the local linear estimation method can be used to estimate the elements of a modified Cholesky decomposition (MCD) of the covariance matrix (Chen and Leng, 2015). The MCD of a covariance matrix **Σ**(*u*) is formulated as

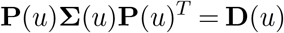

Where 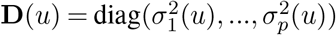, **P**(*u*) is a lower unitriangular matrix with the (*j, l*)-th below diagonal entry being −*ϕ*_*jl*_(*u*), the negative coefficients of the autoregressive models

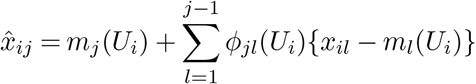

Therefore, the estimators 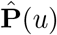 and 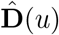 can be obtained by estimating *ϕ*_*jl*_(*u*) and 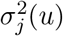, and the covariance estimator 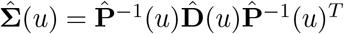 is positive definite. The local linear estimator has been shown to outperform the local constant estimator of the covariance matrix with respect to loss functions in both simulations and data analysis (Chen and Leng, 2015).

Another important direction for the conditional covariance matrix estimation is the expansion to high dimensional covariate (**U**) space. With the plethora of data and computational power available in the modern day, it is inevitable that such estimation would be needed. In nonparametric kernel estimation, the curse of dimensionality is a major problem, where the number of dimensions is so high that the number of samples in a kernel becomes very sparse. There are already nonparametric high-dimension solutions for conditional mean estimation, such as the generalized additive model (GAM; Hastie, 2017). GAM is a nonparametric alternative to linear models, utilizing an iterative backfitting algorithm to estimate the smooth function for each covariate. The additive model avoids the curse of dimensionality by reducing the dimensions via the additive assumption that the covariates are independent. A similar approach to the estimation of the conditional covariance matrix is desired to solve the curse of dimensionality in the covariates space. While there have been studies done in similar topics, there has not been a direct attempt to tackle this problem to the best of my knowledge. For example, a model has been developed for high dimensions in the predictor (**X**) space using MCD (Zhang, 2013). Additionally, a quasi-likelihood based GAM approach has been used to estimate univariate conditional variance functions (Yu, 2019). Yet another manuscript uses MCD of the precision matrix to fit a multivariate model via GAM (Gioia et al., 2022). However, the focus is on outcome estimation while the components of the precision matrix are calculated numerically via model selection.

### 5.2. Conclusion

The methods discussed in this work are ground-breaking for innovation in covariance matrix comparison and application to functional conditional covariance estimation. On one hand, the local linear regression estimator is a well-studied and reliable method capable of performing in multiple situations. On the other hand, the NW kernel estimator for covariance matrices is novel and provides the framework of covariance estimation for different covariate types, such as univariate, bivariate, and mixed data types. Functional covariance matrix estimation has the potential for many types of data and analysis, such as time series, functional data, and graphical modeling. We explore its application to conditional group comparisons in this work.

The three proposed methods for comparing covariance matrices have unique advantages for different situations. The Tracy-Widom method is parametric and can achieve excellent results with a comparatively smaller sample size, given the regularity conditions are met. It strikes a good balance between type I error and power in simulation studies. The permutation method as a nonparametric method has great estimation results given enough sample size, as shown by the type I and power results in the univariate simulations. However, the results worsen in multi-dimensional cases, a well-documented problem for nonparametric estimation (Ramdas et al., 2014). Finally, the Forkman’s test method is powerful but can struggle with high type I error. By combining the ideas of functional conditional covariance estimation and covariance group comparisons, we created a process that can extract a significant amount of information and reach new conclusions that are not feasible with traditional methods.

The simulation studies provided valuable insights into the accuracy of the covariance estimation and the details on the type I error and power of comparison methods. The conditional covariance estimation successfully estimates the true first eigenvalue when the two groups are very different but appears to overestimate when two groups have the same distribution. The strengths and downsides of each comparison method were also explored. For example, the Tracy-Widom method is consistent overall but struggles with low power when the variables are highly correlated. Furthermore, the permutation method can accurately estimate the type I error but struggles with low power, especially in multi-covariate scenarios.

The ADNI biomarkers on dementia data analysis have proven that functional covariance matrix comparison has real-life practical uses in clinical settings. Our method allowed for the comparisons for any given values of age and in auxiliary studies for sex and years of education as well. We gathered more insights by modeling the covariance matrices as a function of the covariates instead of removing these effects. We showed that the covariance matrices of CSF biomarkers are significantly different for younger people but not older ones. However, this was not the case when comparing only female participants. In addition, people with fewer years of education can have significantly different covariance matrices even when they are older. The data analysis clearly demonstrated the utility of functional covariance matrix comparison methods, providing pinpoint results for any given demographics.

While the explored functional covariance comparison methods are innovative and have many benefits, some drawbacks should not be ignored. The NW kernel estimator is subject to boundary bias and, therefore, has less than desirable accuracy at the boundary of the data’s range (Cheruiyot, 2020). Other local estimators, such as local linear in theory, could reduce the boundary bias. However, such estimators could produce covariance matrices that are not positive semi-definite without additional workarounds (Chen and Leng, 2015). In addition, the leave-one-out cross-validation resulted in an optimized covariance bandwidth without great results in type I error and power in simulations. The aforementioned cross-validation criterion minimizes the likelihood separately for each of the two groups, which could be improved upon by using another criterion that considers the likelihood of both groups simultaneously. In addition, other bandwidth optimizers could be used instead, such as a plug-in bandwidth based on quadratic loss or the Stein loss (Fan and Gijbels, 1996). Other bandwidth options could also be considered, such as a kernel contrast method, Do-validation, and bootstrap methods (Heidenreich, Schindler and Sperlich, 2013). Finally, the comparison methods described in this paper require a standardization of the variables, thus becoming a comparison of correlation matrices. Other comparison methods are needed if the magnitude of the variances is important to a study.

## Acknowledgement

Data collection and sharing for this project was funded by the Alzheimer’s Disease Neuroimaging Initiative (ADNI) (National Institutes of Health Grant U01 AG024904) and DOD ADNI (Department of Defense award number W81XWH-12-2-0012). ADNI is funded by the National Institute on Aging, the National Institute of Biomedical Imaging and Bioengineering, and through generous contributions from the following: AbbVie, Alzheimer’s Association; Alzheimer’s Drug Discovery Foundation; Araclon Biotech; BioClinica, Inc.; Biogen; Bristol-Myers Squibb Company; CereSpir, Inc.; Cogstate; Eisai Inc.; Elan Pharmaceuticals, Inc.; Eli Lilly and Company; EuroImmun; F. Hoffmann-La Roche Ltd and its affiliated company Genentech, Inc.; Fujirebio; GE Healthcare; IXICO Ltd.; Janssen Alzheimer Immunotherapy Research & Development, LLC.; Johnson & Johnson Pharmaceutical Research & Development LLC.; Lumosity; Lundbeck; Merck & Co., Inc.; Meso Scale Diagnostics, LLC.; NeuroRx Research; Neurotrack Technologies; Novartis Pharmaceuticals Corporation; Pfizer Inc.; Piramal Imaging; Servier; Takeda Pharmaceutical Company; and Transition Therapeutics. The Canadian Institutes of Health Research is providing funds to support ADNI clinical sites in Canada. Private sector contributions are facilitated by the Foundation for the National Institutes of Health (www.fnih.org). The grantee organization is the Northern California Institute for Research and Education, and the study is coordinated by the Alzheimer’s Therapeutic Research Institute at the University of Southern California. ADNI data are disseminated by the Laboratory for Neuro Imaging at the University of Southern California.

